# Understanding the variation in wood densities of trees and its implications for carbon assessments

**DOI:** 10.1101/523480

**Authors:** Karthik Teegalapalli, Chandan Kumar Pandey, Anand M Osuri, Jayashree Ratnam, Mahesh Sankaran

## Abstract

Wood density is a key functional trait used to estimate aboveground biomass (AGB) and carbon stocks. A common practice in forest AGB and carbon estimation is to substitute genus averages (across species with known wood densities) in cases where wood densities of particular species are unknown. However, the extent to which genus-level averages are reflective of species wood densities across tree genera is uncertain, and understanding this is critical for estimating the accuracy of carbon stock estimates. Using primary field data from India and secondary data from a published global dataset, we quantified the extent to which wood density varied among individuals within species (intraspecific variation) at the regional scale and among species within genera (interspecific variation) at regional to global scales. We used a published global database with wood density data for 7743 species belonging to 1741 genera. Linear models were used to compare the species values with the genera averages and the individual values with the species averages, respectively. To estimate the error associated with using genus-level averages for carbon stocks estimation, we compared genus values averaged at the global, old world and continental scales with species values from actually measured data. We also ran a simulation using vegetation data from a published database to calculate the estimation errors in a 1 hectare plot level when genera-averaged wood densities are used. Intraspecific variation was significantly lower than interspecific variation. Continental level genera averages led to estimates closer to the species values for the 10 genera for which most data on species was available. This was also evident from a comparison of genera averages at these three spatial scales with species values from our data. Species within certain ‘hypervarying’ genera showed relatively high levels of variation, irrespective of the spatial scale of the dataset used. The error in estimation of AGB when genera-averaged values were used for species wood densities was 0.35, 0.71 and 2.43% when 0, 10 and 25% of the girth of the trees in the simulated plot were from hypervariable genera. Our findings indicate that species values provide the most accurate estimates for individuals. Genus average wood density values at the continental scale provided more reliable estimates than those at larger spatial scales. The aboveground biomass estimation error when species wood densities were approximated to the genera-average values was 1.4 to 3.7 tonnes per ha when 10% and 25%, respectively, of the girth of trees was from species from hypervariable genera. Our findings indicate that regional or continental scale genera averages provide more reliable estimates than global data and we propose a method to identify hypervariable genera, for which species values rather than genera averages can provide better estimates of carbon stocks.

## Introduction

Forests provide a critical ecosystem service as carbon repositories, storing more than half of the total carbon stocks in terrestrial ecosystems (Lewis et al. 2009, Pan et al. 2011, Saatchi et al. 2011). With an estimated 20 million hectares of forests lost annually, it is critical to understand the ecosystem value and carbon sequestration potential of the remaining forests (Hansen et al. 2013). Reliable estimation of carbon stocks can be arrived at by incorporating tree wood density, the ratio of the dry mass to the green volume of the wood, in allometric equations used to estimate aboveground biomass (Chave et al. 2005). However, wood densities of most species in a region of interest are not readily available, and genera- or family-level averages are often used instead, since wood density has typically been considered a taxonomically well-conserved trait (Baker et al. 2004, Chave et al. 2006, Flores & Coomes 2011). However, there may be significant phylogenetic and spatial variability in wood density values, therefore, using genus level averages for data-deficient species can potentially lead to significant errors in the estimation of carbon stocks (Chave et al. 2006, Flores & Coomes 2011).

Chave et al. (2006) found that 74% of the variation in species wood density was explained at the genus level and suggest that genus averages provide reliable approximations for species values, except for some hypervariable genera. Flores and Coomes (2011) found that species level approximation using global datasets (*e.g*. Zanne et al. 2009) provide more robust values of wood density than local level data sets, because of larger sample sizes of each genera and species in global databases. To verify this, we quantified variation in wood density among species within genera from continental to global scales to explore the extent of error involved in the estimation of carbon stocks using genera-averaged wood density values at these different scales. Further, acknowledging that wood density values averaged for certain hypervariable genera can lead to inaccurate carbon stock estimates, we sought to provide a method to identify such genera. Our specific research questions were: 1) What is the extent of intraspecific variation in wood density at the regional scale? 2) What is the extent of interspecific variation in wood density at the global, continental and regional spatial scales? and, 3) What is the extent of error associated with using genera averages from global, continental and regional databases, in comparison with actual values?

## Methods

We used a global dataset compiled by Zanne et al. (2009), hereafter, GWWD, and a primary dataset collected by ourselves. For our dataset, we censused 1047 trees from 7 relatively intact forest sites in South India between the years 2012 and 2015 and measured diameter, tree height and wood density according to the methods outlined by Cornelissen et al. (2003). Our final dataset included wood density estimates for 26422 individuals of 7743 species in 1741 genera and 194 families. All analyses were performed using R version 3.4.4 (R Development Core Team, 2018). We used the ‘tpl’ function (Taxonstand package) in R to match synonyms to accepted names for each species and the ‘Plant List’ (http://www.theplantlist.org/) was used as the standard for all species (Cayuela et al. 2012). We retained data with wood density values of four or more species per genera to quantify interspecific variation. To quantify intraspecific variation in wood density, we used the primary data with wood density values of four or more individuals per species.

### Intra- and inter-specific variation in wood density

We used linear models with the primary dataset to compare the wood density of a randomly selected individual with the species average excluding the selected individual to arrive at the intraspecific variation and estimated the mean, variance and R^2^. To understand interspecific variation, we used linear models to compare the wood density of a randomly selected species with its genera average calculated excluding the value of the selected species. This analysis was undertaken with the species values as the response and the genus average as the predictor and repeated at the global, continental and regional scales. To avoid bias due to large sample size, we randomly picked the same number of genera from the global and the continental dataset as were available in the regional dataset and estimated the mean, variance and R^2^ for 500 iterations and compared these across different spatial scales. To identify hypervariable genera, we quantified the number of species with wood density values that were more than 10% of the genus average for the 220 genera for which data for 4 or more species was available. We considered a genus as hypervariable if more than two-thirds of the species within the genus had wood density values greater or less than 10% of the genus average.

### Error in estimation of carbon stocks when using genera averages

Since aboveground biomass is strongly correlated with wood density (Fearnside 1997, Reyes et al. 1992), the ratio of actually measured wood density values and those from genera averages at different spatial scales represents the extent of error associated with using genus averages as substitutes for species values. We also calculated the ratio of species wood density values and genera values from secondary data for the ten most speciose genera within five continents: Asia, Africa, Australia, North America and South and Central America.

### Simulating a 1 hectare plot

We used data from Ramesh et al. (2010) to simulate a 1 ha plot using information on tree girths and species composition. Data was available for approximately 62,000 individuals of which we used a subset of the information on trees and retained the data for which wood density data was available from GWWD and our data. For the simulation, acknowledging that the girth values can contribute to variation in estimated aboveground biomass, we controlled for the overall girth of the simulated 1 ha plot. We randomly selected 1200 trees (girth > 10 cm at breast height), which was approximately the number of trees in our 1 hectare plots in peninsular India and calculated the overall girth which we fixed as the total girth of the simulated plot (Marthews et al. 2015). Using the same criteria as mentioned in the article before, we included 0, 10 and 25% of the overall girth of the plot from hypervariable genera and the rest from non-hypervariable genera in our simulated plot. We chose 10 and 25% of the girth from hypervariable genera since based on our criterion, about 20% of the 220 genera used for our analysis from GWWD were hypervariable (Table S1). Aboveground biomass was calculated using Chave et al. (2005) equation for moist forest stands:

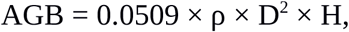

where AGB is the aboveground biomass, ρ is the wood density, D is the diameter of the tree and H is the tree height. We ran the above simulation for 1000 iterations and calculated the percentage difference between aboveground biomass calculated from species and genera value wood densities for the three conditions. We used ANOVA to compare the values from the 3 conditions on a log scale.

## Results

Wood density of the species retained in the dataset for the analysis varied from 0.1 to 1.28 g/cm^3^. A total of 470, 274 and 220 genera had 4 or more species at the global, continental and regional scales, respectively. The results of the linear models with the global, old world and Asian datasets indicated that there was a strong relationship between species and genera values (R^2^> 0.58), the R-square values marginally differed and the slopes and intercepts were comparable at different spatial scales (Figure 1, Table 1). However, for the 10 most speciose genera, the continental level genera averages led to more accurate approximations of species values than global genera averages (Figures S1-S5). Intraspecific variation was significantly lower with a higher R-square value than interspecific variation (R^2^ for intra and interspecific variation was 0.79 and 0.62, respectively, which indicated that species values provided the most accurate values for individuals (Figure 2).

**Table 1:**
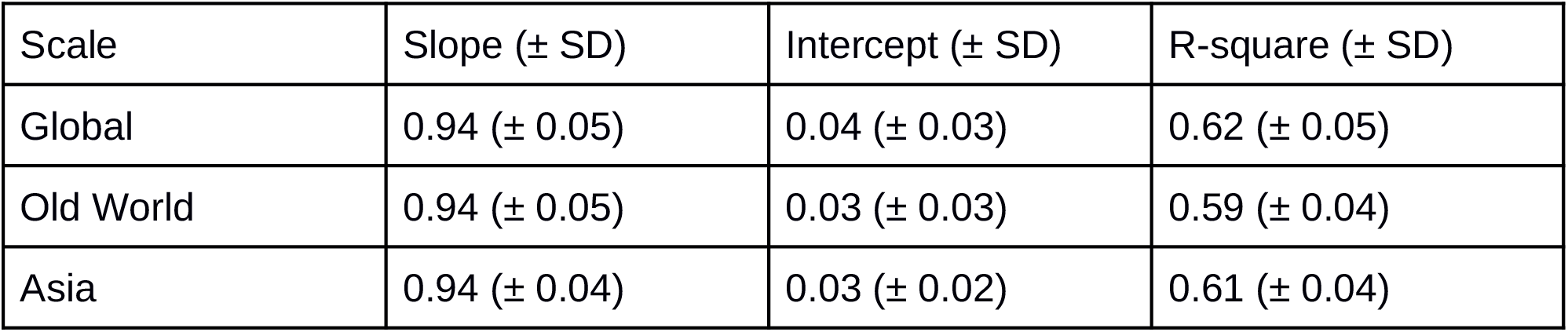
Results of the linear models used to compare the wood density of a randomly selected species with its genera average calculated without the selected species at the global, continental and regional scales for 220 genera.

**Figure 1:**
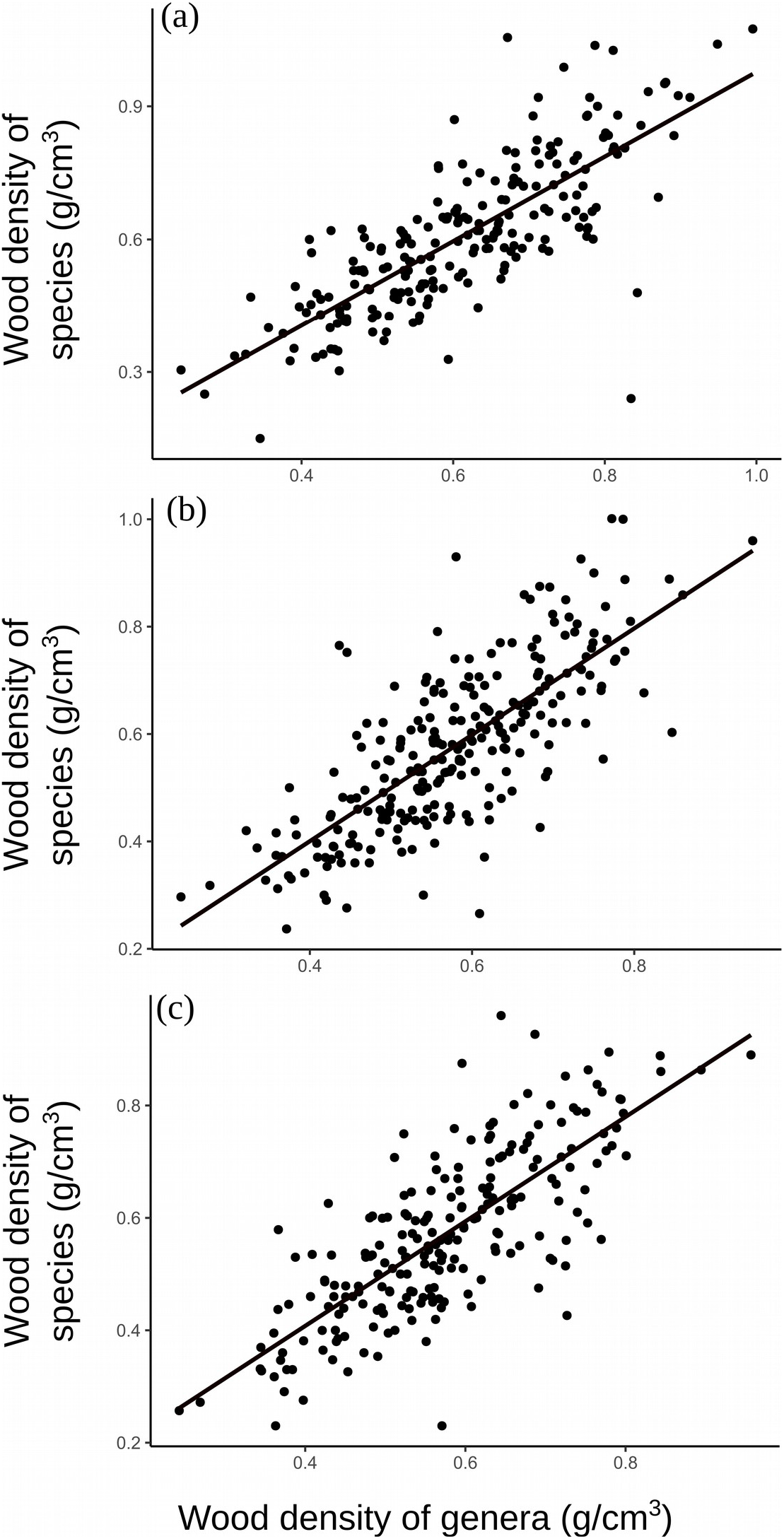
Plot of a linear model with global (a), old world region (b) and Asian (c) datasets with 220 genera that had contained at least 4 species. On the y-axis are the wood density values (g/cm^3^) of randomly selected species and on the x-axis are the genera mean values with the exclusion of the wood density of the the randomly selected species. The R-squared (± SD) values are comparable: 0.623 (± 0.05), 0.587 (± 0.04) and 0.608 (± 0.04) for the global, old world region and Asian datasets indicating that the three datasets provide similar genera approximation for species values.

**Figure 2:**
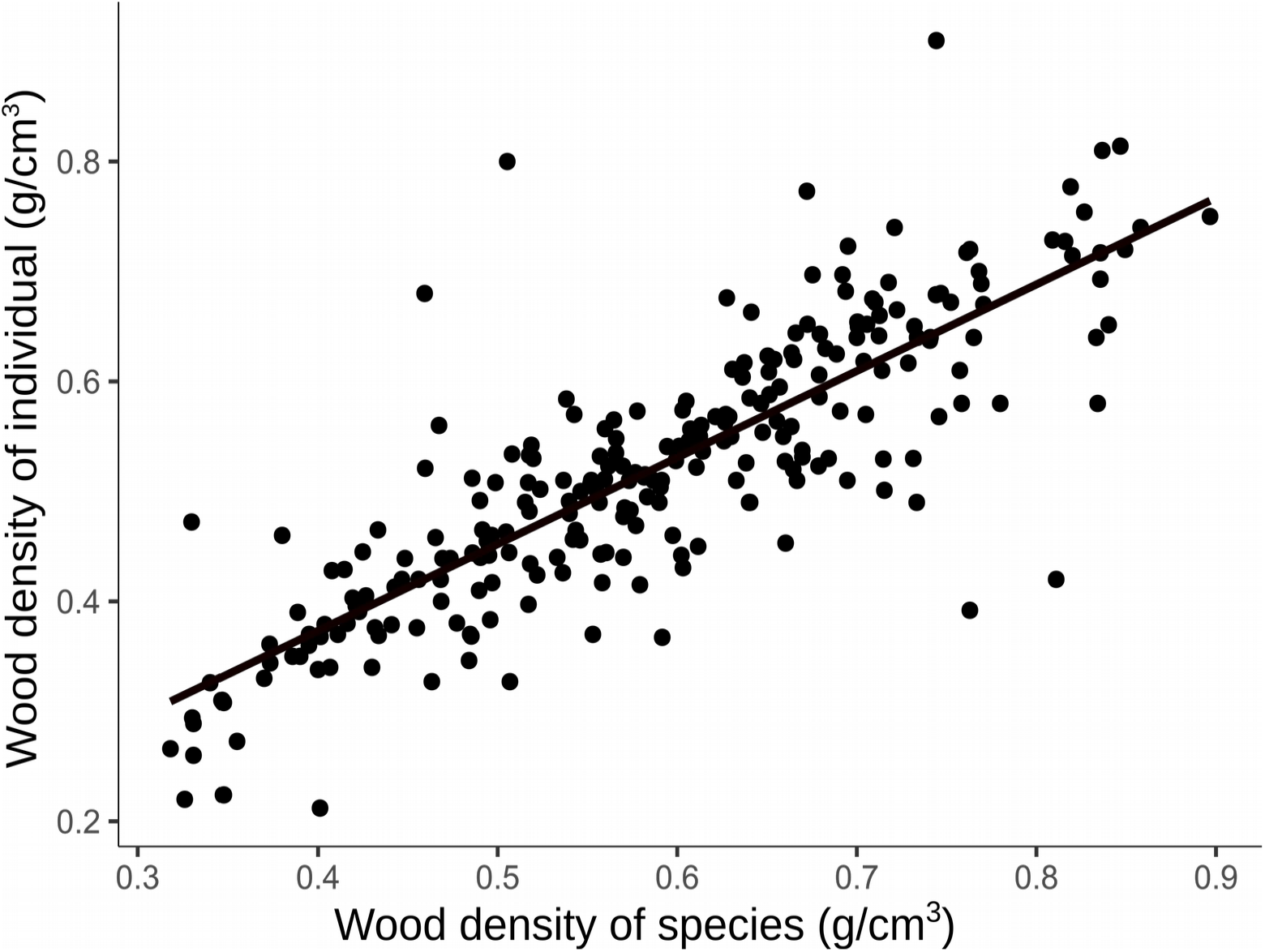
Plot of a linear model with primary data from India with species that had 4 or more individuals within (R^2^: 0.786, N: 132, *p-value:* 1.34e-45). On the y-axis are the wood density values (g/cm^3^) of randomly selected individuals within each species and on the x-axis are the species-level mean values with the exclusion of the wood density of the the randomly selected individual.

### Hypervariable genera

Analysis of the primary data indicated that for species within certain genera such as *Albizia, Dalbergia, Ficus* and *Terminalia*, there was over- or under estimation of wood density irrespective of the scale of the secondary data used for the species-genera approximation (Figure 3). This trend of over- or under estimation of wood density when genera averages were used for species values was observed with the global database as well indicating the hyper-variance of some genera (Figures S1-S5). This was also evident when we tested the number of species within each genus that lie within 10% of the genus average wood density (Table S1). Genera such as *Albizia, Dalbergia, Ficus* and *Terminalia* had several species (53, 61, 59 and 63% of the species, respectively) with wood densities beyond 10% of the genus average. Results from the simulation of a 1 hectare plot indicated an error of 0.35, 0.71 and 2.43% in the calculation of aboveground biomass when genera-averaged wood density values were used for species values, when 0, 10 and 25% of the overall tree girth in the plot was from hypervariable genera (Table 2). For the 1 hectare plot, when the height of the trees is assumed to be 10 m, this indicates an error of (mean ± SE) 0.79 (± 0.02), 1.39 ± (0.03) and 3.7 ± (0.03) tonnes per ha for 0, 10 and 25% of the overall girth from hypervariable genera trees, respectively. ANOVA results indicated a significant positive effect of increasing tree girth from species of hypervariable genera on the aboveground biomass estimation error.

**Figure 3:**
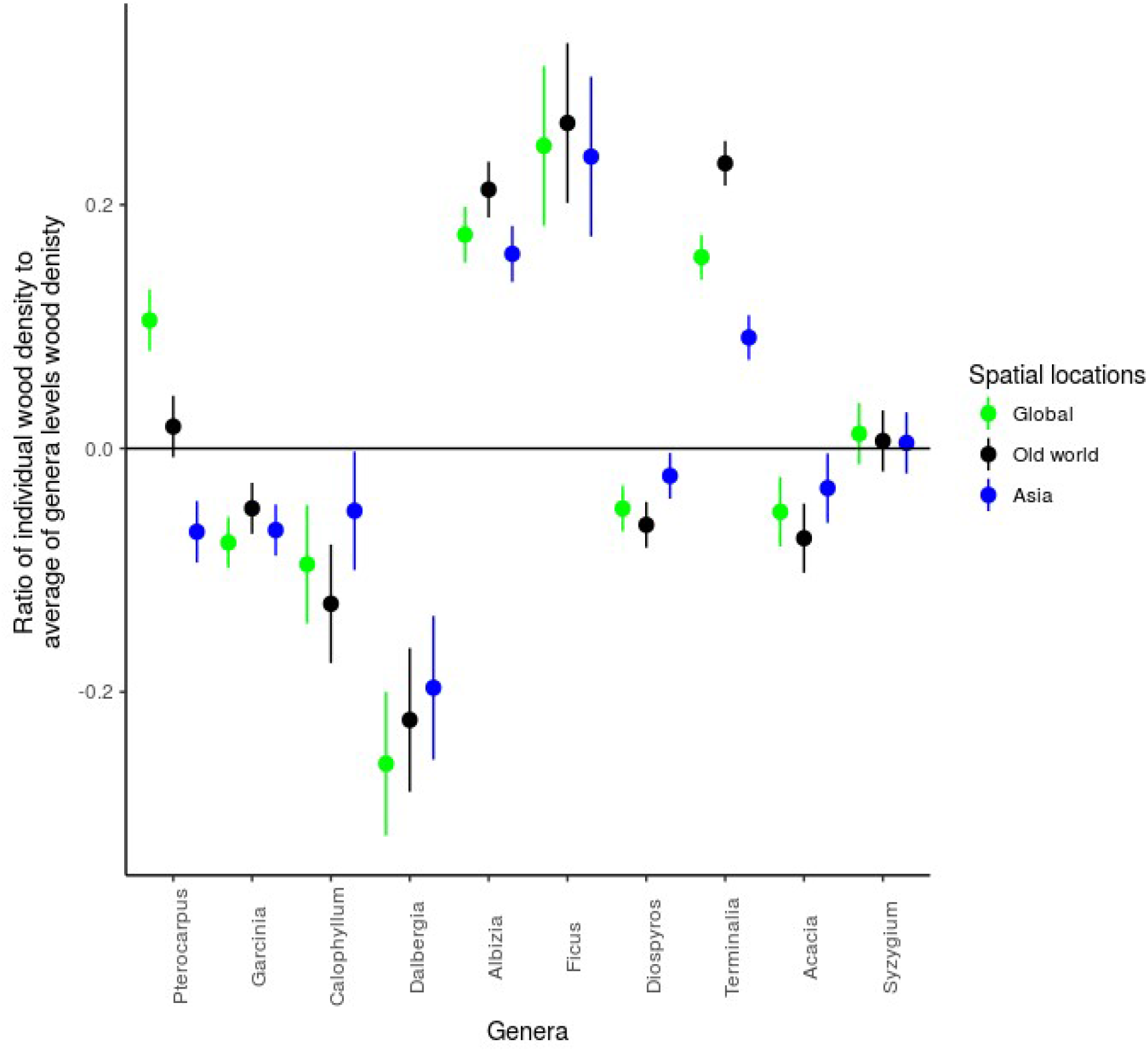
Plot depicting the logarithm of the ratio of individual wood density from primary data to the genera-mean values estimated from global, old world and Asian datasets. The plot indicates that for some genera, wood density based on genera values can be overestimated (ratio < 0) or underestimated (ratio > 0) irrespective of the spatial dataset used for the genera values. This is the case for genera such as *Albizia, Dalbergia, Ficus* and *Terminalia*.

**Table 2:**
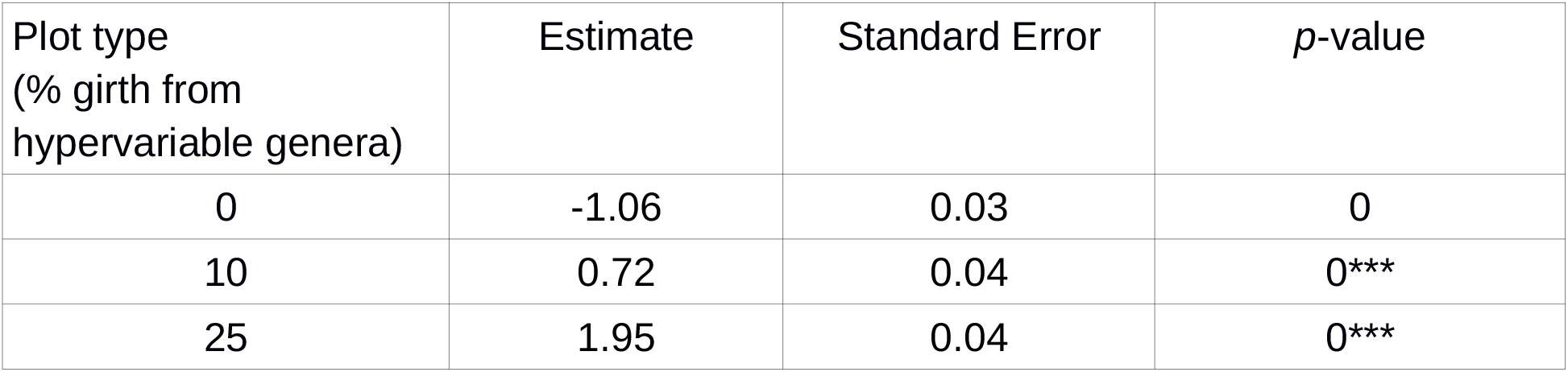
Results of the ANOVA undertaken for aboveground estimation error for a simulated 1 ha plot when genera-averaged wood density values were used for species values. The three conditions under which the error estimates are reported are for 0, 10 and 25% of the overall girth of the trees in the plots from species from hypervariable genera (genera for which more than two-thirds of the species within the genus had wood density values greater or less than 10% of the genus average, *** indicates *p* < 0.001).

| Plot type<br>(% girth from<br>hypervariable genera) | Estimate | Standard Error | $p$-value |
| --- | --- | --- | --- |
| 0 | -1.06 | 0.03 | 0 |
| 10 | 0.72 | 0.04 | 0*** |
| 25 | 1.95 | 0.04 | 0*** |

## Discussion

Continental, rather than global level genus averages, provided more accurate species level approximation on most continents for the top 10 most speciose genera. Contrasting this, Flores & Coomes (2011) suggested that genera level wood densities from global databases provide more accurate species level estimates than regional data sets. However, they also acknowledge that this advantage diminishes with increasing size of the regional dataset. The ratio of species actual wood density to the genera average indicated that for certain genera, irrespective of whether genera-level average was estimated from global or regional values, the values were either over- or under estimated. Analysis of our primary data indicated that this was the case with at least four genera: wood densities of genera such as *Albizia, Ficus* and *Terminalia* were underestimated while *Dalbergia* was overestimated.

The findings of our analysis with global and local data point to analytical steps to reduce errors in carbon stock estimation from wood density data, which we summarise here. Firstly, if wood density values of the species of interest are available, they can be directly used for estimating aboveground biomass and carbon stocks since intraspecific variation in the wood density is generally low. If species values are not available, then genus level wood density averages can be used if the number of species within the genus is sufficient (>4) and the genus is not highly variable in wood density, failing which we recommend that wood density be measured. We suggest that if more than two-third of the species have wood density values that are ± 10% of the genus average, that genus can be considered hypervariable. Such genera can substantially affect aboveground biomass estimation, for instance, in a 1 ha plot that we simulated using data from Ramesh et al. (2010), we found that an aboveground biomass estimation error of 0.8, 1.4 and 3.7 tonnes per ha ensued when 0, 10 and 25 % of the overall tree girth was contributed by trees from hypervariable genera and the estimation error significantly increased with the proportion of tree girth from hypervariable genera.

Using species level values will provide the most accurate aboveground estimates. Our data suggested a tight correlation between species-level and individual-level wood density values, which is consistent with findings of other studies that report low levels of intraspecific variation in comparison with interspecific variation in wood density (Sungpalee et al. 2009, Plourde et al. 2015). For instance, Sungpalee and others (2009) have shown that while intraspecific variation explained only 20%, interspecific variation explained 80% of the total variation in wood density of the sampled individuals. However, we found few studies (Sungpalee et al. 2009, Ogle et al. 2014, Plourde et al. 2015) that have compared intra- vs interspecific variation in wood density and more research is required to corroborate these patterns.

### Conclusions

Our analysis indicated that although genera level approximations for species values were relatively comparable irrespective of the spatial scale of the dataset used, local genera values provided more reliable estimates than global values for the most speciose genera. A low level of intraspecific variability in wood density indicated that species values provide the most accurate estimates for individuals. For certain hypervarying genera, whether local or global genera approximations are used for species, there can be significant under- or overestimation of wood density. To avoid such errors, we have suggested a protocol that can be used to inspect patterns in wood density databases and in specific cases, it may be practical to collect primary data than use secondary data. Overall, our findings with the analysis of wood density databases and the steps outlined to reduce species wood density approximation errors can lead to more accurate estimates of carbon stocks.

## Acknowledgments

The first author acknowledges support from the Department of Biotechnology – Research Associateship. We thank Chengappa SK, Siddarth Machado and Nandita Nataraj for help with the field work involved in this research.

## References

Baker, T.R., Phillips, O.L., Malhi, Y., Almeida, S., Arroyo, L., Di Fiore, A., Erwin, T., Killeen, T.J., Laurance, S.G., Laurance, W.F. & Lewis, S.L., 2004. Variation in wood density determines spatial patterns in Amazonian forest biomass. Global Change Biology, 10, 545–562.

Cayuela, L., Granzow de la Cerda, Í., Albuquerque, F.S. & Golicher, D.J. (2012). Taxonstand: An R package for species names standardisation in vegetation databases. Methods in Ecology and Evolution, 3, 1078–1083.

Chave, J., Andalo, C., Brown, S., Cairns, M.A., Chambers, J.Q., Eamus, D., Fölster, H., Fromard, F., Higuchi, N., Kira, T. & Lescure, J.P. (2005). Tree allometry and improved estimation of carbon stocks and balance in tropical forests. Oecologia, 145, 87–99.

Chave, J., Muller-Landau, H.C., Baker, T.R., Easdale, T.A., Steege, H.T. & Webb, C.O. (2006). Regional and phylogenetic variation of wood density across 2456 neotropical tree species. Ecological Applications, 16, 2356–2367.

Cornelissen, J.H.C., Lavorel, S., Garnier, E., Diaz, S., Buchmann, N., Gurvich, D. E., Reich, P.B., Ter Steege, H., Morgan, H.D., Van Der Heijden, M.G.A. & Pausas, J.G. (2003). A handbook of protocols for standardised and easy measurement of plant functional traits worldwide. Australian journal of Botany, 51, 335–380.

Fearnside, P.M. 1997. Wood density for estimating forest biomass in Brazilian Amazon. Forest Ecology & Management, 90, 59–87.

Flores, O. & Coomes, D.A. (2011). Estimating the wood density of species for carbon stock assessments. Methods in Ecology and Evolution, 2, 214–220.

Hansen, M.C., Potapov, P.V., Moore, R., Hancher, M., Turubanova, S., Tyukavina, A., Thau, D., Stehman, S.V., Goetz, S.J., Loveland, T.R. & Kommareddy, A. (2013). High-resolution global maps of 21st-century forest cover change. Science, 342, 850–853.

Lewis, S.L., Lopez-Gonzalez, G., Sonké, B., Affum-Baffoe, K., Baker, T.R., Ojo, L.O., Phillips, O.L., Reitsma, J.M., White, L., Comiskey, J.A. and Ewango, C.E. (2009). Increasing carbon storage in intact African tropical forests. Nature, 457, 1003.

Marthews, T.R., Nelaballi, S., Ratnam, J., & Sankaran, M. (2015). Ecosystem monitoring and forest census studies in South Asia. Current Science, 108, 1779–1881.

Ogle, K., Pathikonda, S., Sartor, K., Lichstein, J.W., Osnas, J.L., & Pacala, S.W. (2014). A model-based meta-analysis for estimating species-specific wood density and identifying potential sources of variation. Journal of Ecology, 102, 194–208.

Pan, Y., Birdsey, R.A., Fang, J. et al. (2011). A large and persistent carbon sink in the world’s forests. Science, 333, 988–993.

Plourde, B.T., Boukili, V.K. & Chazdon, R.L. (2015). Radial changes in wood specific gravity of tropical trees: inter- and intraspecific variation during secondary succession. Functional Ecology, 29, 111–120.

Ramesh, B.R., Venugopal, P.D., Pélissier, R., Patil, S.V., Swaminath, M.H. & Couteron, P., 2010. Mesoscale patterns in the floristic composition of forests in the central Western Ghats of Karnataka, India. Biotropica, 42, 435–443.

Reyes, G., Brown, S., Chapman, J. & Lugo, A.E. (1992). Wood densities of tropical tree species. General Technical Report SO-88. USDA Forest Service,Southern Forest Experiment Station, New Orleans, Louisiana, USA, 15p.

Saatchi, S.S., Harris, N.L., Brown, S., Lefsky, M., Mitchard, E.T., Salas, W., Zutta, B.R., Buermann, W., Lewis, S.L., Hagen, S. & Petrova, S., 2011. Benchmark map of forest carbon stocks in tropical regions across three continents. Proceedings of the National Academy of Sciences, 108, 9899–9904.

Sungpalee, W., Itoh, A., Kanzaki, M., Sri-ngernyuang, K., Noguchi, H., Mizuno, T., Teejuntuk, S., Hara, M., Chai-udom, K., Ohkubo, T. & Sahunalu, P., (2009). Intra-and interspecific variation in wood density and fine-scale spatial distribution of stand-level wood density in a northern Thai tropical montane forest. Journal of Tropical Ecology, 25, 359–370.

Zanne, A.E., Lopez-Gonzalez, G., Coomes, D.A., Ilic, J., Jansen, S., Lewis, S.L., Miller, R.B., Swenson, N.G., Wiemann, M.C. & Chave, J. (2009). Data from: Towards a worldwide wood economics spectrum. Dryad Digital Repository. https://doi.org/10.5061/dryad.234.

