## Supplemental information for "Understanding the variation in wood densities of trees and its implications for carbon assessments"

**Supplementary information**

*Figures*

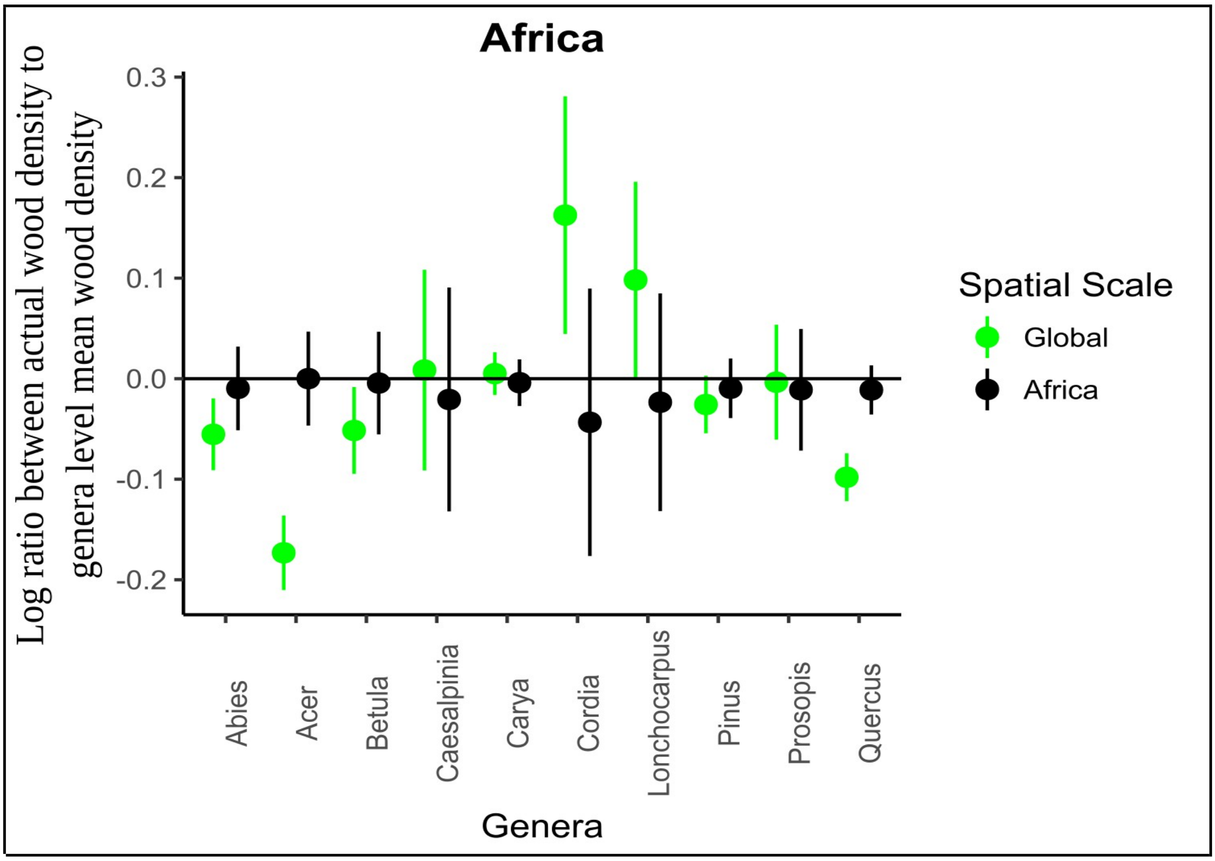


**Figure S1**: The above figure compares the logarithm of the ratio of values of species wood density to the genera average estimated from global data and data only from the Africa continent. The values closer to Y-axis 0 line indicate relatively more accurate approximations of species values from genera averages.

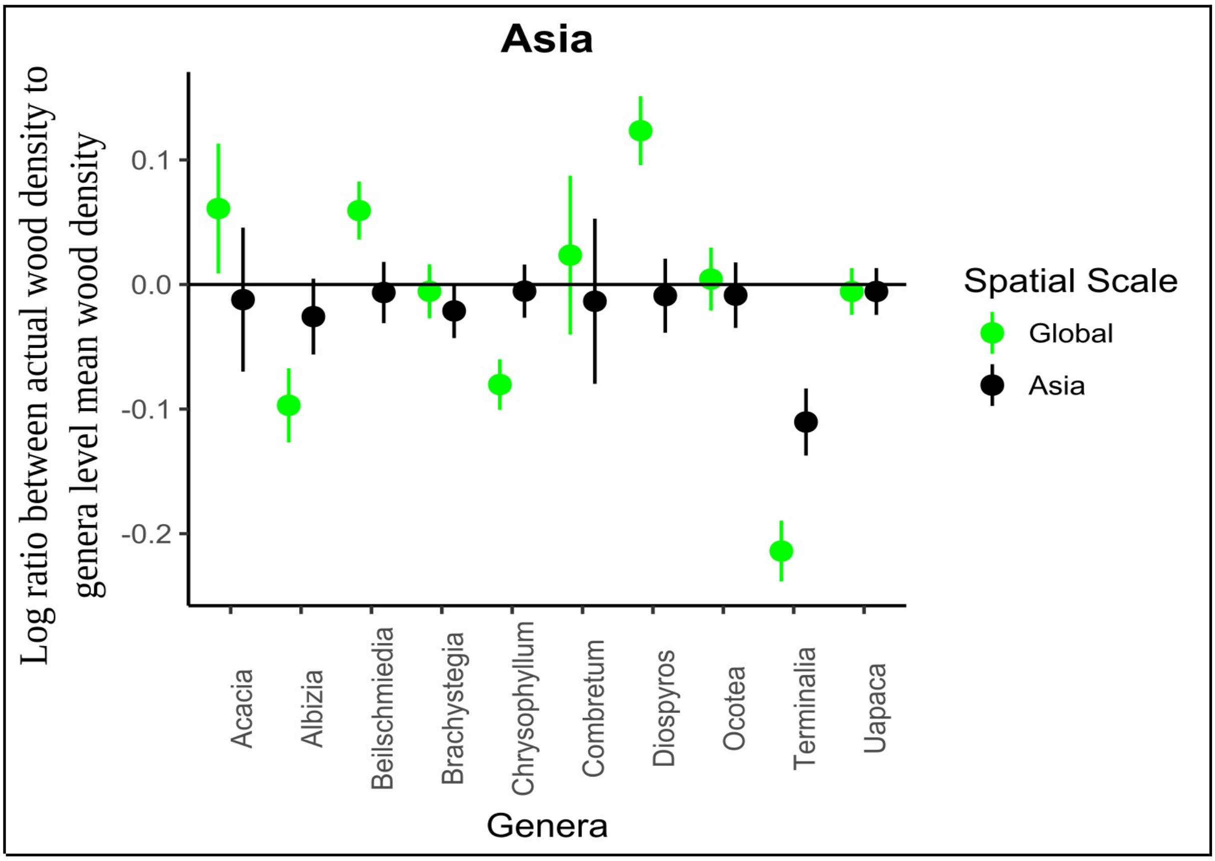


**Figure S2**: The above figure compares the logarithm of the ratio of values of species wood density to the genera average estimated from global data and data only from the Asia continent. The values closer to Y-axis 0 line indicate relatively more accurate approximations of species values from genera averages.

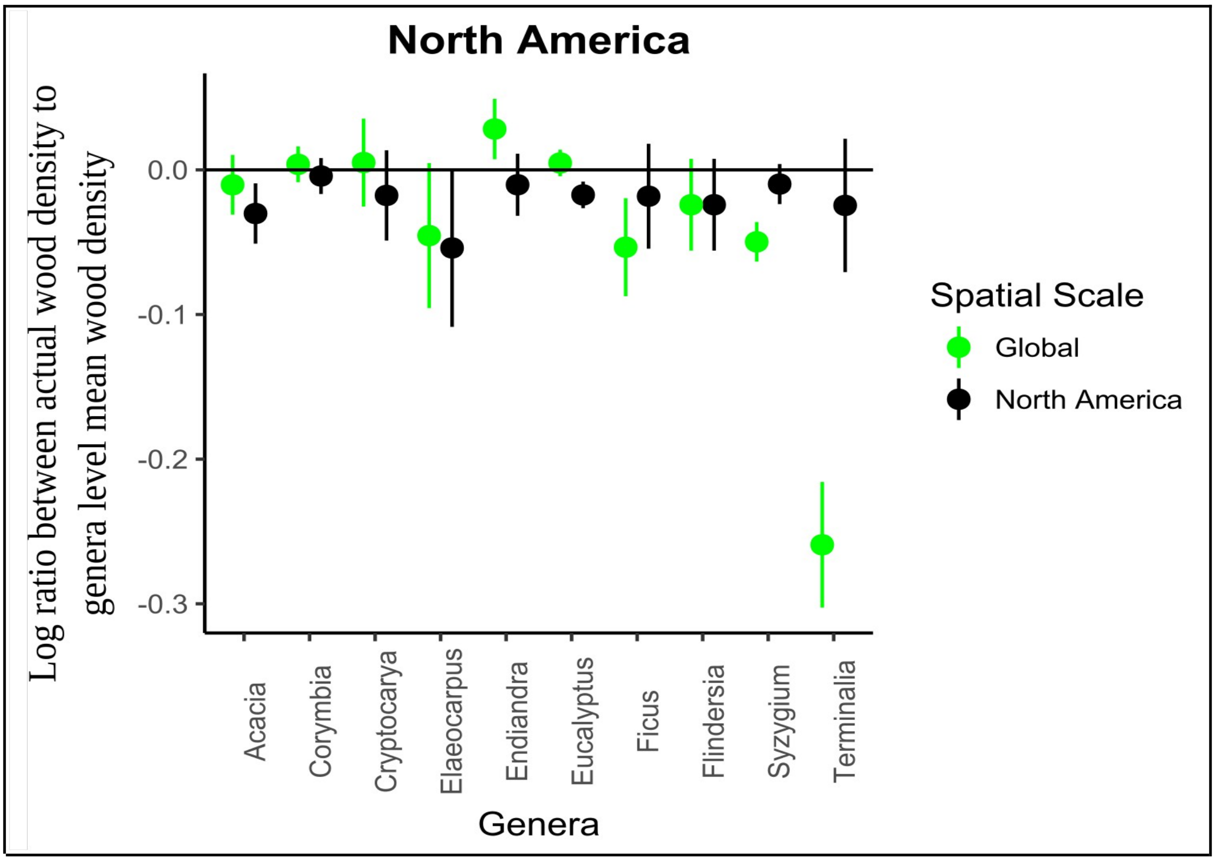


**Figure S3**: The above figure compares the logarithm of the ratio of values of species wood density to the genera average estimated from global data and data only from the North America continent. The values closer to Y-axis 0 line indicate relatively more accurate approximations of species values from genera averages.

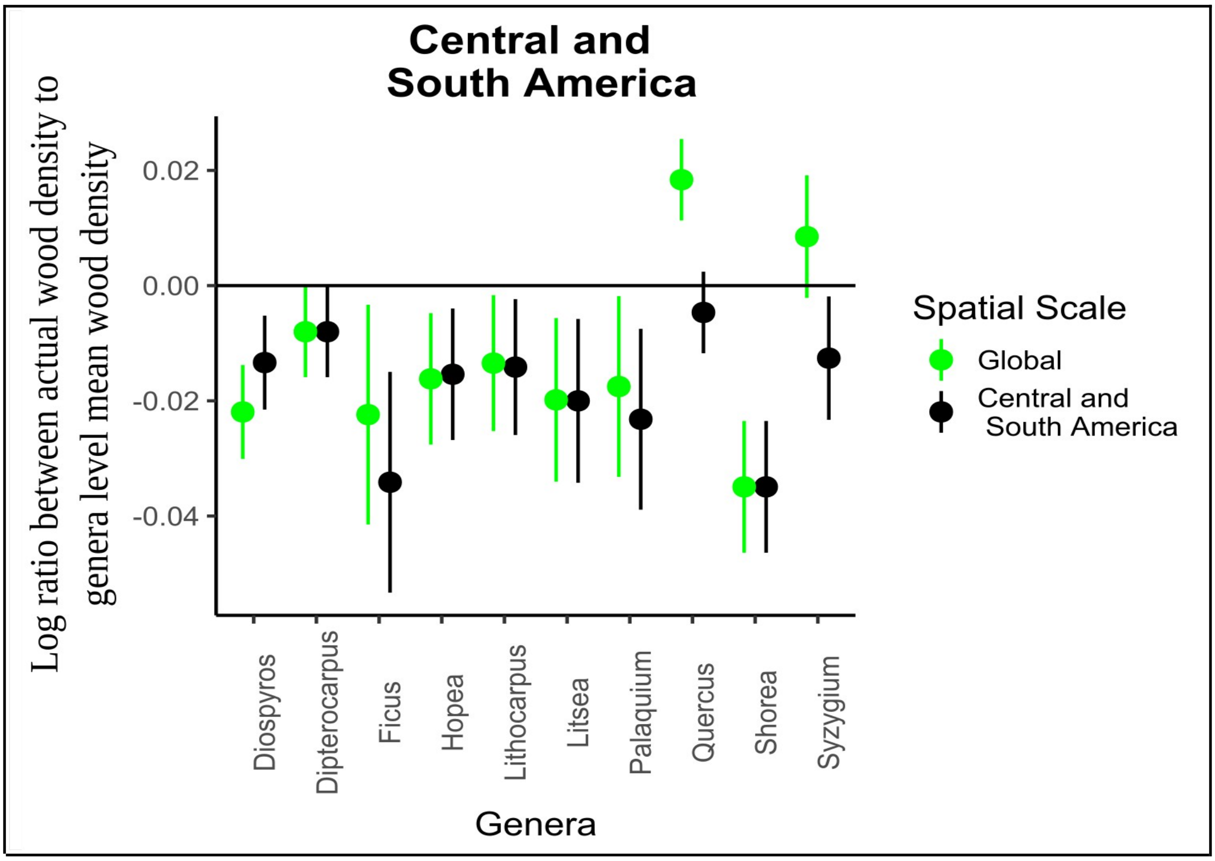


**Figure S4**: The above figure compares the logarithm of the ratio of values of species wood density to the genera average estimated from global data and data only from the South and Central America continent. The values closer to Y-axis 0 line indicate relatively more accurate approximations of species values from genera averages.

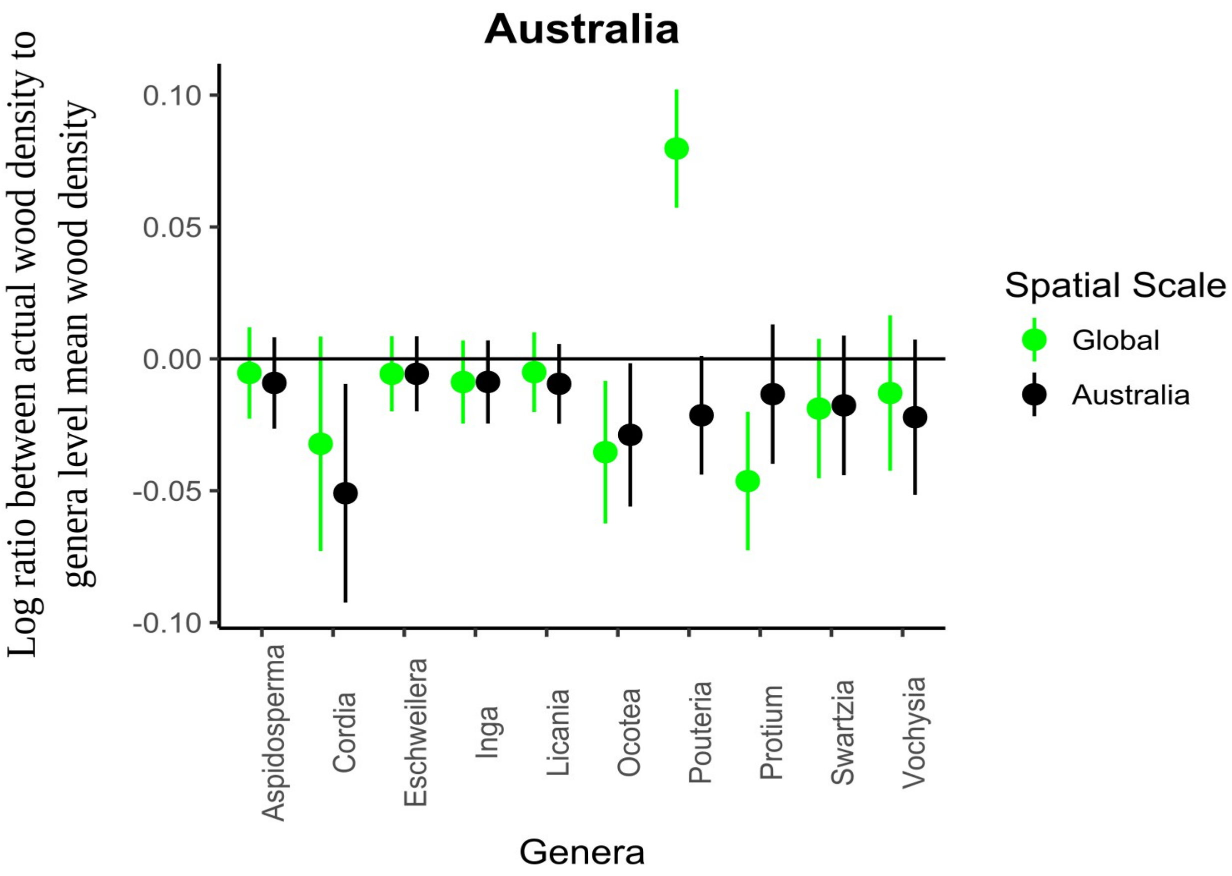


**Figure S5**: The above figure compares the logarithm of the ratio of values of species wood density to the genera average estimated from global data and data only from the Australia continent. The values closer to Y-axis 0 line indicate relatively more accurate approximations of species values from genera averages.

*Table*

**Table S1**: Genera with wood density (WD) data from 4 or more species, the number of species and percentage of species that lie within 10% of the genera mean have been shown.

| Sr. no. | Genus | Number of species within genera | Species with WD beyond 10% genus average WD | % Species with WD beyond 10% genus average WD |
|---|---|---|---|---|
| 1 | *Abies* | 9 | 1 | 11 |
| 2 | *Acacia* | 18 | 12 | 67 |
| 3 | *Acer* | 12 | 7 | 58 |
| 4 | *Acronychia* | 6 | 2 | 33 |
| 5 | *Actinodaphne* | 9 | 6 | 67 |
| 6 | *Adenanthera* | 4 | 2 | 50 |
| 7 | *Adinandra* | 18 | 6 | 33 |
| 8 | *Agathis* | 8 | 3 | 38 |
| 9 | *Aglaia* | 32 | 12 | 38 |
| 10 | *Ailanthus* | 4 | 3 | 75 |
| 11 | *Alangium* | 11 | 9 | 82 |
| 12 | *Albizia* | 17 | 9 | 53 |
| 13 | *Alnus* | 5 | 1 | 20 |
| 14 | *Alphitonia* | 5 | 2 | 40 |
| 15 | *Alphonsea* | 6 | 2 | 33 |
| 16 | *Alseodaphne* | 12 | 7 | 58 |
| 17 | *Alstonia* | 9 | 7 | 78 |
| 18 | *Altingia* | 4 | 0 | 0 |
| 19 | *Anisophyllea* | 5 | 3 | 60 |
| 20 | *Anisoptera* | 9 | 2 | 22 |
| 21 | *Anogeissus* | 6 | 0 | 0 |
| 22 | *Antidesma* | 9 | 4 | 44 |
| 23 | *Aporosa* | 16 | 7 | 44 |
| 24 | *Archidendron* | 5 | 2 | 40 |
| 25 | *Ardisia* | 6 | 2 | 33 |
| 26 | *Artocarpus* | 30 | 17 | 57 |
| 27 | *Astronia* | 4 | 1 | 25 |
| 28 | *Baccaurea* | 13 | 6 | 46 |
| 29 | *Barringtonia* | 7 | 4 | 57 |
| 30 | *Bauhinia* | 4 | 1 | 25 |
| 31 | *Beilschmiedia* | 17 | 6 | 35 |
| 32 | *Betula* | 8 | 2 | 25 |
| 33 | *Bombax* | 5 | 0 | 0 |
| 34 | *Bridelia* | 7 | 4 | 57 |
| 35 | *Bruguiera* | 5 | 0 | 0 |
| 36 | *Buchanania* | 7 | 3 | 43 |
| 37 | *Burckella* | 4 | 0 | 0 |
| 38 | *Caldcluvia* | 4 | 4 | 100 |
| 39 | *Callicarpa* | 4 | 2 | 50 |
| 40 | *Calophyllum* | 31 | 13 | 42 |
| 41 | *Campnosperma* | 4 | 2 | 50 |
| 42 | *Canarium* | 38 | 20 | 53 |
| 43 | *Casearia* | 10 | 2 | 20 |
| 44 | *Castanopsis* | 37 | 25 | 68 |
| 45 | *Celtis* | 11 | 3 | 27 |

| | | | | |
|---|---|---|---|---|
| 46 | *Chionanthus* | 5 | 2 | 40 |
| 47 | *Chisocheton* | 6 | 2 | 33 |
| 48 | *Cinnamomum* | 26 | 10 | 38 |
| 49 | *Citrus* | 4 | 1 | 25 |
| 50 | *Cleistanthus* | 4 | 3 | 75 |
| 51 | *Colona* | 5 | 0 | 0 |
| 52 | *Cordia* | 7 | 6 | 86 |
| 53 | *Cratoxylum* | 5 | 4 | 80 |
| 54 | *Croton* | 8 | 4 | 50 |
| 55 | *Crudia* | 15 | 8 | 53 |
| 56 | *Cryptocarya* | 24 | 8 | 33 |
| 57 | *Cupressus* | 7 | 0 | 0 |
| 58 | *Cynometra* | 7 | 1 | 14 |
| 59 | *Dacrycarpus* | 5 | 2 | 40 |
| 60 | *Dacrydium* | 4 | 1 | 25 |
| 61 | *Dacryodes* | 7 | 3 | 43 |
| 62 | *Dalbergia* | 18 | 11 | 61 |
| 63 | *Daphniphyllum* | 4 | 0 | 0 |
| 64 | *Dehaasia* | 6 | 3 | 50 |
| 65 | *Dialium* | 8 | 4 | 50 |
| 66 | *Dillenia* | 21 | 8 | 38 |
| 67 | *Diospyros* | 93 | 44 | 47 |
| 68 | *Dipterocarpus* | 47 | 21 | 45 |
| 69 | *Dolichandrone* | 4 | 2 | 50 |
| 70 | *Drepananthus* | 5 | 2 | 40 |
| 71 | *Dryobalanops* | 7 | 2 | 29 |
| 72 | *Drypetes* | 15 | 6 | 40 |
| 73 | *Durio* | 17 | 3 | 18 |
| 74 | *Dysoxylum* | 22 | 14 | 64 |
| 75 | *Elaeocarpus* | 27 | 11 | 41 |
| 76 | *Endiandra* | 8 | 4 | 50 |
| 77 | *Endospermum* | 7 | 3 | 43 |
| 78 | *Erythrina* | 5 | 2 | 40 |
| 79 | *Erythroxylum* | 4 | 0 | 0 |
| 80 | *Eucalyptus* | 11 | 8 | 73 |
| 81 | *Euodia* | 5 | 3 | 60 |
| 82 | *Eurya* | 4 | 2 | 50 |
| 83 | *Fagraea* | 8 | 5 | 63 |
| 84 | *Ficus* | 54 | 32 | 59 |
| 85 | *Flacourtia* | 5 | 1 | 20 |
| 86 | *Fraxinus* | 8 | 2 | 25 |
| 87 | *Garcinia* | 40 | 26 | 65 |
| 88 | *Gardenia* | 7 | 2 | 29 |
| 89 | *Glochidion* | 12 | 4 | 33 |
| 90 | *Gluta* | 15 | 8 | 53 |
| 91 | *Gmelina* | 6 | 2 | 33 |
| 92 | *Gonystylus* | 13 | 4 | 31 |
| 93 | *Gordonia* | 6 | 1 | 17 |
| 94 | *Grewia* | 11 | 5 | 45 |
| 95 | *Gymnacranthera* | 6 | 3 | 50 |
| 96 | *Haplolobus* | 5 | 5 | 100 |

| | | | | |
|---|---|---|---|---|
| 97 | *Helicia* | 6 | 3 | 50 |
| 98 | *Heritiera* | 11 | 5 | 45 |
| 99 | *Hibiscus* | 6 | 5 | 83 |
| 100 | *Homalium* | 10 | 2 | 20 |
| 101 | *Hopea* | 62 | 35 | 56 |
| 102 | *Horsfieldia* | 13 | 6 | 46 |
| 103 | *Hydnocarpus* | 15 | 7 | 47 |
| 104 | *Ilex* | 9 | 6 | 67 |
| 105 | *Ixonanthes* | 6 | 2 | 33 |
| 106 | *Ixora* | 7 | 2 | 29 |
| 107 | *Kayea* | 6 | 3 | 50 |
| 108 | *Knema* | 18 | 5 | 28 |
| 109 | *Lagerstroemia* | 16 | 4 | 25 |
| 110 | *Larix* | 6 | 3 | 50 |
| 111 | *Lepisanthes* | 4 | 2 | 50 |
| 112 | *Lithocarpus* | 44 | 22 | 50 |
| 113 | *Litsea* | 44 | 30 | 68 |
| 114 | *Lophopetalum* | 14 | 11 | 79 |
| 115 | *Macaranga* | 33 | 26 | 79 |
| 116 | *Machilus* | 7 | 2 | 29 |
| 117 | *Madhuca* | 33 | 18 | 55 |
| 118 | *Magnolia* | 28 | 16 | 57 |
| 119 | *Mallotus* | 13 | 7 | 54 |
| 120 | *Mangifera* | 17 | 7 | 41 |
| 121 | *Manilkara* | 8 | 1 | 13 |
| 122 | *Maniltoa* | 4 | 2 | 50 |
| 123 | *Mastixia* | 5 | 4 | 80 |
| 124 | *Melanochyla* | 8 | 5 | 63 |
| 125 | *Melicope* | 7 | 1 | 14 |
| 126 | *Meliosma* | 9 | 7 | 78 |
| 127 | *Memecylon* | 18 | 8 | 44 |
| 128 | *Mesua* | 8 | 7 | 88 |
| 129 | *Microcos* | 12 | 9 | 75 |
| 130 | *Miliusa* | 6 | 1 | 17 |
| 131 | *Millettia* | 4 | 4 | 100 |
| 132 | *Mischocarpus* | 4 | 0 | 0 |
| 133 | *Mitragyna* | 4 | 1 | 25 |
| 134 | *Mitrephora* | 6 | 2 | 33 |
| 135 | *Morus* | 7 | 2 | 29 |
| 136 | *Myristica* | 27 | 13 | 48 |
| 137 | *Nauclea* | 4 | 3 | 75 |
| 138 | *Neolitsea* | 6 | 5 | 83 |
| 139 | *Neonauclea* | 13 | 4 | 31 |
| 140 | *Nephelium* | 10 | 5 | 50 |
| 141 | *Nothofagus* | 6 | 4 | 67 |
| 142 | *Ochrosia* | 4 | 2 | 50 |
| 143 | *Olea* | 7 | 2 | 29 |
| 144 | *Ormosia* | 12 | 8 | 67 |
| 145 | *Palaquium* | 48 | 30 | 63 |
| 146 | *Parartocarpus* | 4 | 2 | 50 |
| 147 | *Parashorea* | 9 | 5 | 56 |

| | | | | |
|---|---|---|---|---|
| 148 | *Parinari* | 5 | 1 | 20 |
| 149 | *Parishia* | 6 | 1 | 17 |
| 150 | *Parkia* | 6 | 4 | 67 |
| 151 | *Paulownia* | 5 | 1 | 20 |
| 152 | *Payena* | 10 | 6 | 60 |
| 153 | *Pentace* | 9 | 4 | 44 |
| 154 | *Persea* | 7 | 2 | 29 |
| 155 | *Phoebe* | 10 | 3 | 30 |
| 156 | *Phyllanthus* | 4 | 1 | 25 |
| 157 | *Picea* | 8 | 2 | 25 |
| 158 | *Pinus* | 25 | 10 | 40 |
| 159 | *Pittosporum* | 5 | 0 | 0 |
| 160 | *Planchonella* | 12 | 9 | 75 |
| 161 | *Planchonia* | 5 | 2 | 40 |
| 162 | *Podocarpus* | 8 | 2 | 25 |
| 163 | *Polyalthia* | 16 | 11 | 69 |
| 164 | *Polyscias* | 4 | 1 | 25 |
| 165 | *Populus* | 16 | 9 | 56 |
| 166 | *Pouteria* | 6 | 4 | 67 |
| 167 | *Premna* | 4 | 2 | 50 |
| 168 | *Prunus* | 18 | 11 | 61 |
| 169 | *Pternandra* | 6 | 3 | 50 |
| 170 | *Pterocarpus* | 7 | 3 | 43 |
| 171 | *Pterocymbium* | 5 | 3 | 60 |
| 172 | *Pterospermum* | 10 | 7 | 70 |
| 173 | *Ptychopyxis* | 4 | 0 | 0 |
| 174 | *Quercus* | 42 | 10 | 24 |
| 175 | *Radermachera* | 5 | 4 | 80 |
| 176 | *Rhizophora* | 4 | 0 | 0 |
| 177 | *Rhodamnia* | 4 | 0 | 0 |
| 178 | *Rhus* | 4 | 0 | 0 |
| 179 | *Ryparosa* | 9 | 4 | 44 |
| 180 | *Sageraea* | 4 | 2 | 50 |
| 181 | *Salix* | 7 | 5 | 71 |
| 182 | *Sandoricum* | 4 | 1 | 25 |
| 183 | *Santiria* | 9 | 7 | 78 |
| 184 | *Sapindus* | 5 | 3 | 60 |
| 185 | *Saraca* | 4 | 3 | 75 |
| 186 | *Sarcosperma* | 4 | 0 | 0 |
| 187 | *Saurauia* | 5 | 3 | 60 |
| 188 | *Scaphium* | 5 | 1 | 20 |
| 189 | *Schima* | 4 | 0 | 0 |
| 190 | *Schizomeria* | 5 | 1 | 20 |
| 191 | *Semecarpus* | 4 | 4 | 100 |
| 192 | *Shorea* | 114 | 90 | 79 |
| 193 | *Sindora* | 17 | 3 | 18 |
| 194 | *Sloanea* | 13 | 6 | 46 |
| 195 | *Sonneratia* | 5 | 3 | 60 |
| 196 | *Stemonurus* | 4 | 0 | 0 |
| 197 | *Sterculia* | 15 | 12 | 80 |
| 198 | *Stereospermum* | 6 | 2 | 33 |

| | | | | |
|---|---|---|---|---|
| 199 | *Streblus* | 6 | 4 | 67 |
| 200 | *Strombosia* | 5 | 3 | 60 |
| 201 | *Strychnos* | 4 | 0 | 0 |
| 202 | *Styrax* | 5 | 3 | 60 |
| 203 | *Swintonia* | 7 | 1 | 14 |
| 204 | *Symplocos* | 7 | 4 | 57 |
| 205 | *Syzygium* | 61 | 33 | 54 |
| 206 | *Tabernaemontana* | 5 | 2 | 40 |
| 207 | *Taxus* | 4 | 4 | 100 |
| 208 | *Teijsmanniodendron* | 6 | 5 | 83 |
| 209 | *Terminalia* | 27 | 17 | 63 |
| 210 | *Ternstroemia* | 4 | 1 | 25 |
| 211 | *Tilia* | 7 | 4 | 57 |
| 212 | *Toona* | 4 | 2 | 50 |
| 213 | *Tristaniopsis* | 4 | 0 | 0 |
| 214 | *Turpinia* | 4 | 2 | 50 |
| 215 | *Ulmus* | 8 | 5 | 63 |
| 216 | *Vatica* | 19 | 9 | 47 |
| 217 | *Vitex* | 11 | 8 | 73 |
| 218 | *Wrightia* | 4 | 1 | 25 |
| 219 | *Xanthophyllum* | 13 | 4 | 31 |
| 220 | *Xylopia* | 5 | 4 | 80 |
